# Decoding personalized motor cortical excitability states from human electroencephalography

**DOI:** 10.1101/2021.10.22.465447

**Authors:** Sara J Hussain, Romain Quentin

## Abstract

Brain state-dependent transcranial magnetic stimulation (TMS) requires real-time identification of cortical excitability states. Current approaches deliver TMS during brain states that correlate with motor cortex (M1) excitability at the group level. Here, we hypothesized that machine learning classifiers could successfully discriminate between high and low M1 excitability states in individual participants using information obtained from low-density electroencephalography (EEG) signals. To test this, we analyzed a publicly available dataset that delivered 600 single TMS pulses to the right M1 during EEG and electromyography (EMG) recordings in 20 healthy adults. Multivariate pattern classification was used to discriminate between brain states during which TMS evoked small and large motor-evoked potentials (MEPs). Results show that personalized classifiers successfully discriminated between low and high M1 excitability states in 80% of tested participants. MEPs elicited during classifier-predicted high excitability states were significantly larger than those elicited during classifier-predicted low excitability states in 90% of tested participants. Personalized classifiers did not generalize across participants. Overall, results show that individual participants exhibit unique brain activity patterns which predict low and high M1 excitability states and that these patterns can be efficiently captured using low-density EEG signals. Our findings suggest that deploying individualized classifiers during brain state-dependent TMS may enable fully personalized neuromodulation in the future.

## 1. Introduction

Over the past few decades, transcranial magnetic stimulation (TMS) has emerged as a potential treatment for a variety of brain disorders, including depression,^1,2^ substance use disorders,^3–5^ memory problems,^6,7^ and motor impairment caused by neurological damage.^8–10^ Although early work reported that different patterns of repetitive TMS significantly alter primary motor cortex (M1) excitability,^11–14^ such interventions have since been shown to exhibit weak effect sizes and wide inter- and intra-individual variability,^15,16^ with in some cases less than half of tested individuals showing the expected plastic response.^17^

Brain state-dependent TMS may improve the magnitude and consistency of TMS effects by delivering TMS during periods of increased cortical excitability and thus increased sensitivity to LTP-like plasticity.^18–22^ In the motor system, brain state-dependent TMS typically involves applying stimulation during brain states that correlate with M1 excitability at the group level, like sensorimotor mu rhythm desynchronization^18,19,23^ or sensorimotor mu rhythm trough phases.^20–23^ Results from studies using this approach have so far been promising: M1 TMS interventions applied during high excitability brain states potentiate corticospinal excitability^18–21^ and enhance motor memory^22^ more than identical interventions applied during low excitability states.

Nonetheless, current brain state-dependent TMS approaches pose some challenges. Most notably, a reliable high excitability brain state must first be detected across a range of individuals. This can be difficult, as shown by several recent studies reporting inconsistent relationships between the sensorimotor mu rhythm and corticospinal excitability.^20,24–27^ Further, maximizing the beneficial effects of brain state-dependent TMS requires the targeted brain state to positively correlate with excitability in each individual. Excluding individuals who do not show such a correlation is also problematic, as it limits the generalizability of results and may artificially inflate effect sizes by screening out potential “non-responders” ahead of time.^26^ Other obstacles include difficulty extracting the targeted brain state in all individuals,^28^ a problem that will only be exacerbated in clinical studies involving patients with brain lesions^29^ or other forms of volumetric brain loss.

Here, we developed an analytical approach that overcomes these issues by exploiting the rich spatiotemporal dynamics embedded within whole-scalp EEG signals. Specifically, we aimed to identify personalized patterns of brain activity that predict the presence of high and low M1 excitability states. We hypothesized that multivariate pattern classification, a form of supervised machine learning, could successfully discriminate between high and low M1 excitability states in individual participants using only simple power spectral features calculated from low-density EEG signals. To test our hypothesis, we trained personalized machine learning classifiers to identify brain states during which single-pulse TMS evoked either large or small motor-evoked potentials (MEPs) in 20 healthy adults. Our results show that these personalized classifiers successfully discriminated between high and low M1 excitability states in 80% of tested participants. Further, personalized classifiers did not generalize across participants, indicating that individual participants exhibit unique brain activity patterns that predict high and low M1 excitability states. Overall, our findings provide the conceptual and methodological framework for efficient, fully personalized decoding-based brain state-dependent TMS interventions in the future.

## 2. Materials and Methods

### 2.1. Human participants

Data analyzed in this study have been published previously^25,30^ and are publicly available (https://openneuro.org/datasets/ds002094/versions/1.0.0). The original study involved simultaneous single-pulse transcranial magnetic stimulation (TMS), electroencephalography (EEG), and electromyography (EMG) in 20 healthy participants (6 F and 14 M, age 30 ± 1.59 [SEM] years). Original study procedures were approved by the National Institutes of Health Combined Neuroscience Section Institutional Review Board and conformed to local regulations for human subjects research. All participants provided their written informed consent prior to participation. The original study involved a 5-minute eyes-open resting EEG recording followed by application of 600 single TMS pulses during EEG and EMG recordings (see^25^ for additional methodological details). Only EEG and EMG data acquired during TMS application are presented here.

### 2.2. Data acquisition

#### 2.2.1. EEG and EMG signal acquisition

EEG signals were recorded using TMS-compatible amplifiers (BrainAmp MR+, BrainVision; sampling rate = 5 kHz, hardware filtering = DC – 1 kHz, resolution 0.5 μV). Bipolar EMG signals were recorded using adhesive electrodes arranged in a belly-tendon montage (Signal, Cambridge Electronic Design; sampling rate = 5 kHz, hardware filtering = 5 Hz – 2 kHz).

#### 2.2.2. TMS

TMS was applied as previously reported.^25,30^ Briefly, monophasic posterior-to-anterior current TMS (MagStim 200^2^, MagStim Co. Ltd) was applied over the scalp hotspot for the left first dorsal interosseous muscle (L. FDI) using a figure-of-eight coil. After determining resting motor threshold (RMT) using an automatic threshold-tracking algorithm (Adaptive PEST procedure^31^), 6 blocks of 100 single TMS pulses were applied to the L. FDI hotspot at 120% RMT (inter-stimulus interval = 5 s with 15% jitter). Breaks were taken during each block when requested by the subject or in response to TMS coil overheating. To minimize fatigue throughout the experiment, participants were offered breaks after each block of 100 pulses and encouraged to request extra breaks within each block when desired. TMS coil position was monitored for accuracy using frameless neuronavigation throughout the experiment (BrainSight, Rogue Research).

### 2.3. Data and statistical analysis

#### 2.3.1. EMG processing

EMG signals acquired during single-pulse TMS delivery were used to calculate peak-to-peak MEP amplitudes, which served as an objective marker of M1 excitability. EMG data were analyzed using custom-written scripts in MATLAB combined with FieldTrip.^32^ EMG data were segmented from 25 ms to 100 ms after TMS delivery, demeaned, and linearly detrended. Residual EMG offset was removed by subtracting the average EMG amplitude between 1 and 20 ms before TMS delivery from the entire timeseries signal. MEPs contaminated by voluntary motor activity were identified as those trials that contained (1) prestimulus EMG root-mean-square values exceeding an individually-defined upper limit (75^th^ percentile + 3 * the inter-quartile range) or (2) MEP signals that were poorly correlated with the average MEP signal observed in that participant (Pearson’s correlation, r < 0.4). These MEPs were excluded from further analysis (7.52 ± 1.40% of all MEPs). In each remaining trial, the peak-to-peak MEP amplitude was calculated as the difference between the maximum and minimum voltage deflection between 20 and 40 ms after each TMS pulse. Low and high M1 excitability states were defined at the individual participant level as those brain states during which single-pulse TMS elicited an MEP with an amplitude in the top 50% or bottom 50% of all remaining MEP amplitudes, respectively (i.e., median split). MEP amplitudes were dichotomized in this way because brain state-dependent TMS requires a binary decision (i.e., whether to deliver TMS) to be made at each time step during real-time signal processing.

#### 2.3.2. EEG preprocessing

EEG signals preceding single-pulse TMS delivery were used to calculate spectral power across a range of frequencies. Resulting spectral power values were used as features when building participant-specific classifiers. EEG data were analyzed using custom-written scripts in MATLAB combined with FieldTrip.^32^ EEG signals acquired during single-pulse TMS delivery were re-referenced to the common average reference and then segmented into six time windows leading up to TMS delivery (windows included 3.005 – 2.505 s, 2.505 – 2.005 s, 2.005 – 1.505 s, 1.505 – 1.005 s, 1.005 – 0.505 s and 0.505 – 0.005 s before TMS). Data from TP9 and TP10 were removed due to poor scalp electrode contact; thus, 28 channels remained for analysis (FP1, FP2, F3, F4, C3, C4, P3, P4, O1, O2, F7, F8, T7, T8, P7, P8, Fz, Cz, Pz, Iz, FC1, FC2, CP1, CP2, FC5, FC6, CP5, and CP6). For each time window, data were demeaned, linearly detrended and downsampled to 500 Hz to reduce computational load. Power spectra were calculated for each time window using Welch’s method (4-100 Hz with 0.25 Hz resolution, excluding line noise frequencies [58-62 Hz]). Power spectral estimates were averaged within each frequency band separately for each window. This created 168 power spectral features per window (i.e., 28 EEG channels for each of 6 different frequency bands). Features included broadband (4-100 Hz, excluding line noise frequencies [58-62 Hz]), theta (4-7 Hz), alpha (8-12 Hz), beta (13-35 Hz), low gamma (36-58 Hz), and high gamma (62-100 Hz) power. The frequencies used to define broadband and band-limited power ranges were informed by previous studies.^20,22,25,33–35^

#### 2.3.3. Single-trial optimization and classification at the individual participant level

Multivariate pattern analysis (MVPA) was performed using the MVPA-Light toolbox^36^ combined with custom-written scripts in MATLAB. We aimed to predict whether an MEP elicited by a single TMS pulse was either small or large (see *EMG processing*) using information contained within power spectral features obtained immediately preceding each pulse (0.505 – 0.005 s before TMS; see *EEG preprocessing*). To achieve this, we used participant-specific Linear Discriminant Analysis (LDA) with stratified k-fold cross-validation (k = 5). This allowed each participant’s personalized classifier to find the hyperplane that best separated the low and high M1 excitability states.

We used k-fold cross-validated grid search and feature ranking to optimize two key criteria when building personalized classifiers: 1) the number of power spectral features included in the classifier, and 2) the lambda value (i.e., learning rate). We first divided each participant’s trials into 5 folds. Then, we iteratively searched for both the optimal number of features and the optimal lambda value by training an LDA classifier on 80% of all available trials (i.e., 4 of 5 folds) and then testing this classifier on the remaining 20% of trials (i.e., the remaining fold). For each fold, we optimized the number of features used by first ranking all 168 power spectral features by importance using chi-square tests. That is, we tested for statistical dependencies between each feature and the two M1 excitability states (i.e., high or low, see *EMG preprocessing*) using each fold’s training data (i.e., 80% of all available trials). The negative log of the chi-square test’s p-value was calculated as the feature score; here, higher feature scores reflect features that more strongly covary with M1 excitability states within that fold’s training data. All features were ranked by importance using the feature score. During the grid search, features were added in order of importance (i.e., classifier 1 used the single most important feature, classifier 2 used the two most important features, etc.), such that at each iteration, the LDA classifier used a feature set that had 1 more feature than the previous iteration. Feature ranking was always performed on the training set only. To optimize each participant’s lambda value within each fold, we also trained and tested 100 LDA classifiers for each set of features, with 100 possible lambda values linearly spaced between 1e-10 and 1. In each participant, a total of 16,800 cross-validated LDA classifiers (168 possible feature sets x 100 possible lambda values, with 5 folds per classifier) were trained and used to predict M1 excitability state (high or low) for each trial. For each cross-validated classifier, we calculated the area under the receiver-operator curve (AUC, 1 = perfect classification, 0 = completely inaccurate classification) by comparing each classifier’s predicted MEP classes to the ground truth MEP classes per fold. AUC values were averaged across folds to obtain a performance metric for each of the 16,800 classifiers. Training and testing sets were always independent and stratification was used. Optimal lambda values and number of features were on average 0.08 ± 0.04 and 45.1 ± 11.03, respectively. After grid search, each participant’s best performing cross-validated classifier was used to calculate a confusion matrix indicating the percentage of accurate and inaccurate classifications. Confusion matrices were averaged across participants to obtain a single group-level confusion matrix. See Figure 1 for a schematic illustration of the full optimization and classification pipeline. On average, optimizing each participant’s personalized classifier using this procedure took 4.02 ± 0.05 minutes (range = 3.63 – 4.54) on a standard laptop computer (MacBook Pro, 2.3 GHz Quad-Core Intel i7 processor, 16 GB RAM).

**Figure 1.**
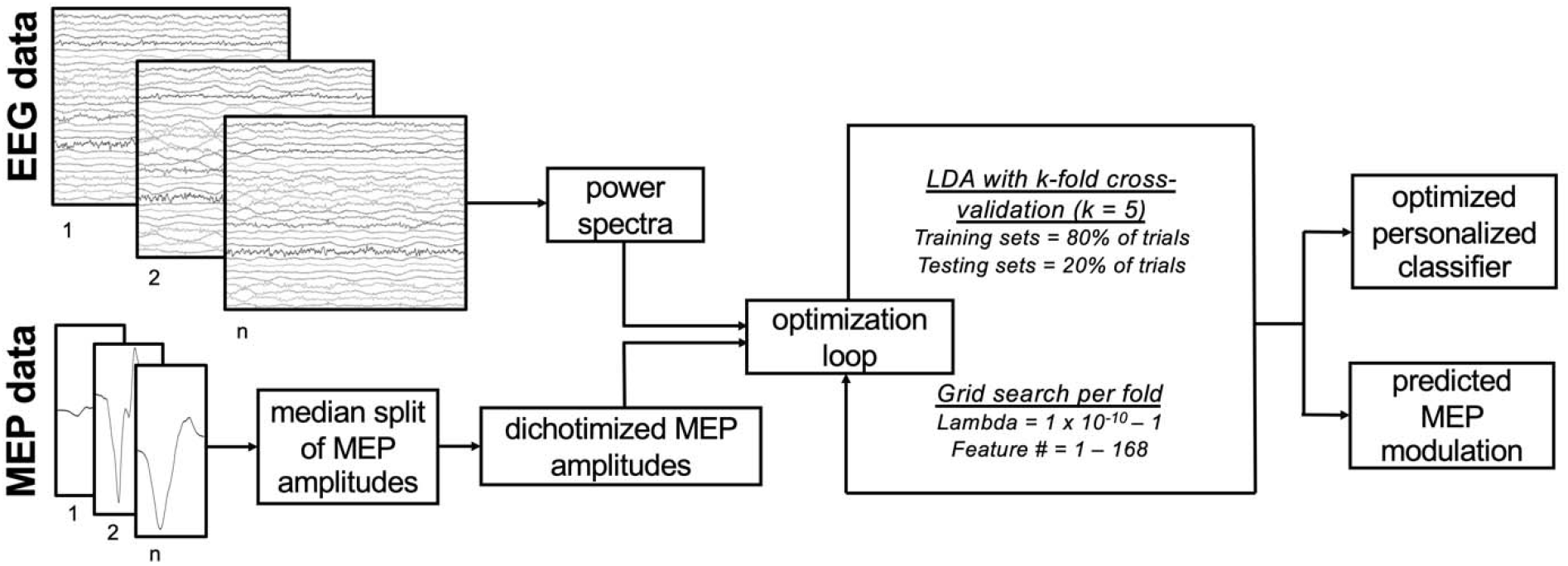
Analytical procedures used to optimize each participant’s personalized classifier. Raw EEG data undergo spectral analysis. MEP amplitudes are calculated and divided into small and large MEPs via median split. Afterwards, 5-fold cross-validated grid search is performed by fitting LDA classifiers for 100 different lambda values and 168 different feature numbers, with features added in order of importance. Then, each participant’s best performing cross-validated classifier is selected and the predicted MEP amplitude modulation is calculated using the true MEP amplitudes for each of the predicted classes obtained during cross-validation. For each participant, 16,800 classifiers are trained and tested during the optimization and classification loop, which takes ~4 minutes on a standard laptop computer.

Performance of each participant’s optimized classifier was statistically evaluated using permutation tests. During permutation tests, the same k-fold cross-validated grid search classification procedure described above was used, except that instead of optimizing classifiers to successfully discriminate between true low and true high M1 excitability states, classifiers were optimized to discriminate between random excitability states, defined as randomly shuffled true M1 excitability states. This cross-validated grid search procedure with permuted excitability states was performed 500 times, producing a distribution of 500 AUC values reflecting the performance of optimized null classifiers for each participant. If a given participant’s true AUC value (as determined during k-fold cross-validated grid search classification, see above) exceeded the 95^th^ percentile of this null distribution (analogous to a right-tailed statistical test with alpha = 0.05), above-chance classification was considered to have occurred. At the group level, null AUC values exhibited a mean of 0.55; all group-level analyses of performance at the individual level therefore used 0.55 as the empirical chance level for our two-class classification problem. Group-level classification performance was thus evaluated statistically by comparing the group-level distribution of true AUC values to this chance level. Other than permutation tests, all statistical testing was performed using right-tailed Wilcoxon signed rank tests. Alpha was equal to 0.05 for all analyses.

To confirm that each participant’s optimized classifier correctly predicted the presence of low and high M1 excitability states, we also compared the amplitudes of MEPs evoked during classifier-predicted low M1 excitability states and classifier-predicted high M1 excitability states at the individual and group levels. This analysis was carried out using predicted MEP classes obtained from the best performing classifier obtained during cross-validated grid search (see above). At the individual level, MEP amplitudes were compared between predicted low and predicted high M1 excitability states using a left-tailed Mann Whitney U-test. The U-test was used instead of the Wilcoxon signed rank test to account for potentially unequal trial numbers per predicted class. For group-level analysis, MEP amplitudes were averaged within each classifier-predicted state per participant, pooled across participants, and then compared using a left-tailed Wilcoxon signed rank test. Left-tailed statistical tests were used to test the *a priori* hypothesis that MEPs elicited during classifier-predicted low excitability states would be smaller than those elicited during classifier-predicted high excitability states. To quantify the maximum possible MEP amplitude difference, we also performed the same analysis steps for MEP amplitudes elicited during true low and true high M1 excitability states separated by the median split (see *EMG processing*). Alpha was equal to 0.05 for all analyses.

#### 2.3.4. Generalization of personalized classifiers across participants

To test the ability of a personalized classifier to correctly classify low and high M1 excitability states in other participants, we trained the previously personalized and optimized LDA classifier (see 2.2.3: *Single trial classification at the individual participant level*) using all trials of the corresponding participant. This trained classifier was then used to discriminate between low and high M1 excitability states in other participants, using all trials obtained from each other participant as separate testing sets. Such generalization across participants was trained and tested using the same time window immediately preceding each pulse (0.505 – 0.005 s before TMS; see *EEG preprocessing*). Generalization performance was assessed using AUC values. Because classifiers were trained on one participant and tested on all others, training and testing sets were always independent. Generalization performance was evaluated at the individual participant and group levels. For individual participants, AUC values reflecting the extent to which each participant’s personalized classifier generalized to all other test participants was compared to theoretical chance level (0.5). At the group level, AUC values were averaged across all 19 test participants per training participant. This created a distribution of aggregate generalization AUC values, which were then also compared to the theoretical chance level (0.5). Finally, aggregate generalization AUC values were compared to AUC values obtained during single-trial classification at the individual level. All statistical testing was performed using right-tailed Wilcoxon signed rank tests. Alpha was equal to 0.05 for all analyses.

#### 2.3.5. Classification and generalization on earlier time windows

To determine if classifiers performed differently when trained and tested using power spectral features obtained from earlier time windows preceding TMS, we applied the same k-fold cross-validated grid search classification approach described above (see *Single-trial optimization and classification at the individual participant level*), except that classifiers were instead trained and tested using power spectral features from EEG signals obtained within five different 500 ms windows preceding TMS (3.005 – 2.505 s, 2.505 – 2.005 s, 2.005 – 1.505 s, 1.505 – 1.005 s, 1.005 – 0.505 s before TMS). Classification performance for each time window was evaluated statistically by comparing group-level AUC values to the empirical chance level for individual classification (0.55) using right-tailed false discovery rate (FDR)-corrected Wilcoxon signed rank tests.^37^ We also used pairwise FDR-corrected Wilcoxon signed rank tests to determine if each time window’s AUC values exceeded the AUC values obtained from classifiers trained using EEG features obtained immediately before TMS delivery (0.505 – 0.005 s before TMS). Because results of this analysis revealed above-chance classification performance at all tested time windows (see *Results*), we further asked whether classifiers trained on the time-window preceding the TMS pulse (0.505 – 0.005 s before TMS) generalized to earlier time windows (3.005 – 2.505 s, 2.505 – 2.005 s, 2.005 – 1.505 s, 1.505 – 1.005 s, 1.005 – 0.505 s before TMS). This was done using the same k-fold cross-validated grid search classification approach described above. AUC values describing the extent to which each participant’s personalized classifier generalized backwards in time were thus calculated.^38^ Group-level temporal generalization performance was evaluated by comparing AUC values at each time window both to the empirical chance level for individual classification (0.55) and to AUC values obtained from the time window preceding the TMS pulse (0.505 – 0.005 s before TMS) using right-tailed FDR-corrected Wilcoxon signed rank tests. Alpha was equal to 0.05 for all analyses.

#### 2.3.6. Influence of trial number on classification performance and classifier-predicted MEP amplitude modulation

We further determined how many trials are needed to achieve successful classification of M1 excitability states. First, we divided each participant’s full dataset into smaller datasets with 50, 100, 150, 200, 250, 300, 350, 400 and 450 trials. 450 trials was chosen as the maximum dataset size because it was the largest increment of 50 trials for which all participants had enough trials available. Trials within each dataset were selected in order of their occurrence (i.e., each participant’s 50-trial dataset contained trials 1 – 50, each participant’s 100-trial dataset contained trials 1 – 100, etc.). We then performed the same k-fold cross-validated grid search classification approach described above (see *Single-trial optimization and classification at the individual participant level*) on each of these smaller datasets. Group-level AUC values were compared to the empirical chance level for individual classification (0.55, see *Single-trial optimization and classification at the individual participant level*) for each dataset size using right-tailed Wilcoxon signed-rank tests and compared between dataset sizes using FDR-corrected two-tailed Wilcoxon signed-rank tests. Group-level classifier-predicted MEP amplitude modulation values were compared to one (i.e., no MEP amplitude modulation) for each dataset size and then compared across dataset sizes using FDR-corrected two-tailed Wilcoxon signed-rank tests. Alpha was equal to 0.05 for all analyses.

#### 2.3.7. Evaluation of computational efficiency

To determine the tractability of our approach in real time, we also calculated the time needed to compute power spectral features and perform classification. For each participant, the time needed to complete these steps was tested on 500 ms data segments obtained from each trial. Computational times were averaged across all data segments per participant. Right-tailed Spearman’s rho was used to determine if the number of features incorporated into each participant’s final classifier correlated with that participant’s mean computational time. Alpha was equal to 0.05.

## 3. Results

We first determined if personalized classifiers trained using features obtained immediately before each TMS pulse (power spectral features calculated from EEG data within 0.505 to 0.005 s before TMS) successfully discriminated between low and high M1 excitability states. 16 of 20 participants showed above-chance classification performance (p < 0.05 for all) while the remaining 4 participants did not (p > 0.11 for all). Similarly, group-level classification performance was significantly higher than empirical chance (chance = 0.55, AUC = 0.62 ± 0.01 [range = 0.53 – 0.72], p < 6.5 × 10^-5^; Figure 2a). The group-level confusion matrix confirmed that accurate classification occurred more often than inaccurate classification (Figure 2b, yellow versus blue cells). MEP amplitudes elicited during classifier-predicted high excitability states were significantly larger than those elicited during classifier-predicted low excitability states in 18 participants (p < 0.02 for all) while for 2 participants this was not the case (p > 0.10 for all, Figure 2c). At the group level, MEP amplitudes were 23.68 ± 2.98% larger (range = −8.86 – 44.12%) when evoked during classifier-predicted high versus classifier-predicted low excitability states (p < 1.36 × 10^-4^, Figure 2c). For reference, MEPs elicited during true high excitability states were significantly larger than those elicited during true low excitability states in all participants (p < 8.08 × 10^-77^ for all) and at the group level (p < 4.78 × 10^-5^, Figure 2d). MEPs elicited during true high excitability states were 258.84 ±22.94% larger (range = 57.17 – 428.5%) than MEPs elicited during true low excitability states. Of all possible power spectral features, gamma features across the scalp were most often included in each participant’s final classifier (Figure 3).

**Figure 2.**
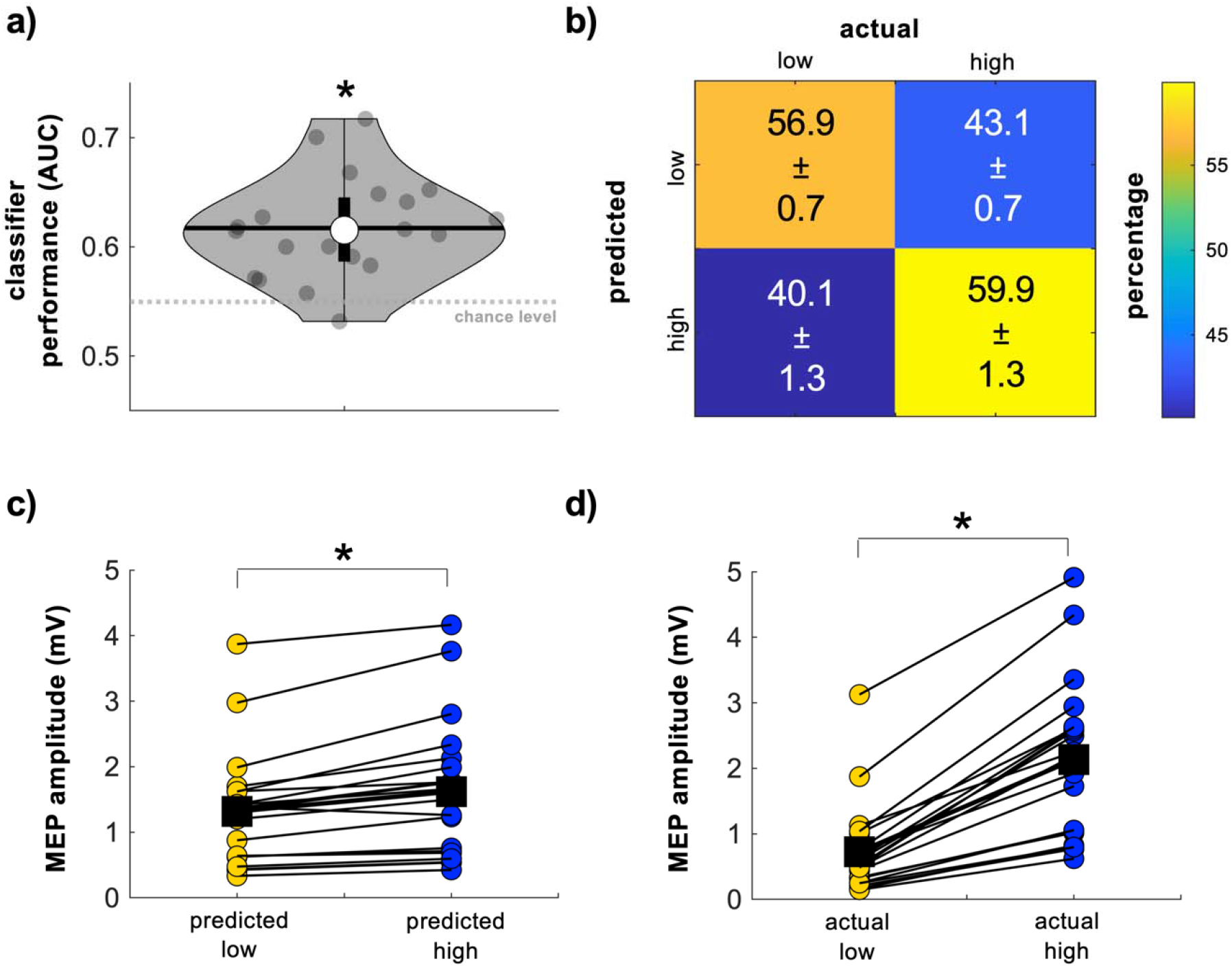
Personalized classification of low and high M1 excitability states. a) AUC values reflecting classification performance. Classifiers were trained at the individual participant level using personalized power spectral features obtained from EEG data immediately preceding TMS (i.e., between 0.505 and 0.005 s before TMS). These personalized classifiers performed significantly above chance in 16 of 20 participants and at the group level. The grey dashed line indicates the empirical chance level for a two-class classification problem (here, 0.55). Grey dots = individual participant AUC values, thick whisker = interquartile range, thin whisker = full range, white dot = group median, black horizontal line = group mean, grey dashed line = empirical chance level. b) Group-level confusion matrix describing the percentage of accurate and inaccurate classifications of M1 excitability states. Note that the percentage of true low (upper left) and true high (lower right) classifications exceed the percentage of false low (upper right) and false high (lower left) classifications. Error estimates are SEM. c) MEP amplitudes elicited during classifier-predicted low (yellow dots) and classifier-predicted high (blue dots) M1 excitability states. MEPs were significantly larger during classifier-predicted high excitability states than classifier-predicted low excitability states in 18 of 20 participants (colored dots) and at the group level (black squares). c) MEP amplitudes elicited during true low (yellow dots) and true high (blue dots) M1 excitability states. As expected, MEPs were significantly larger during true high excitability states than true low excitability states in all participants (colored dots) and at the group level (black squares). * Indicates group-level statistical significance.

**Figure 3.**
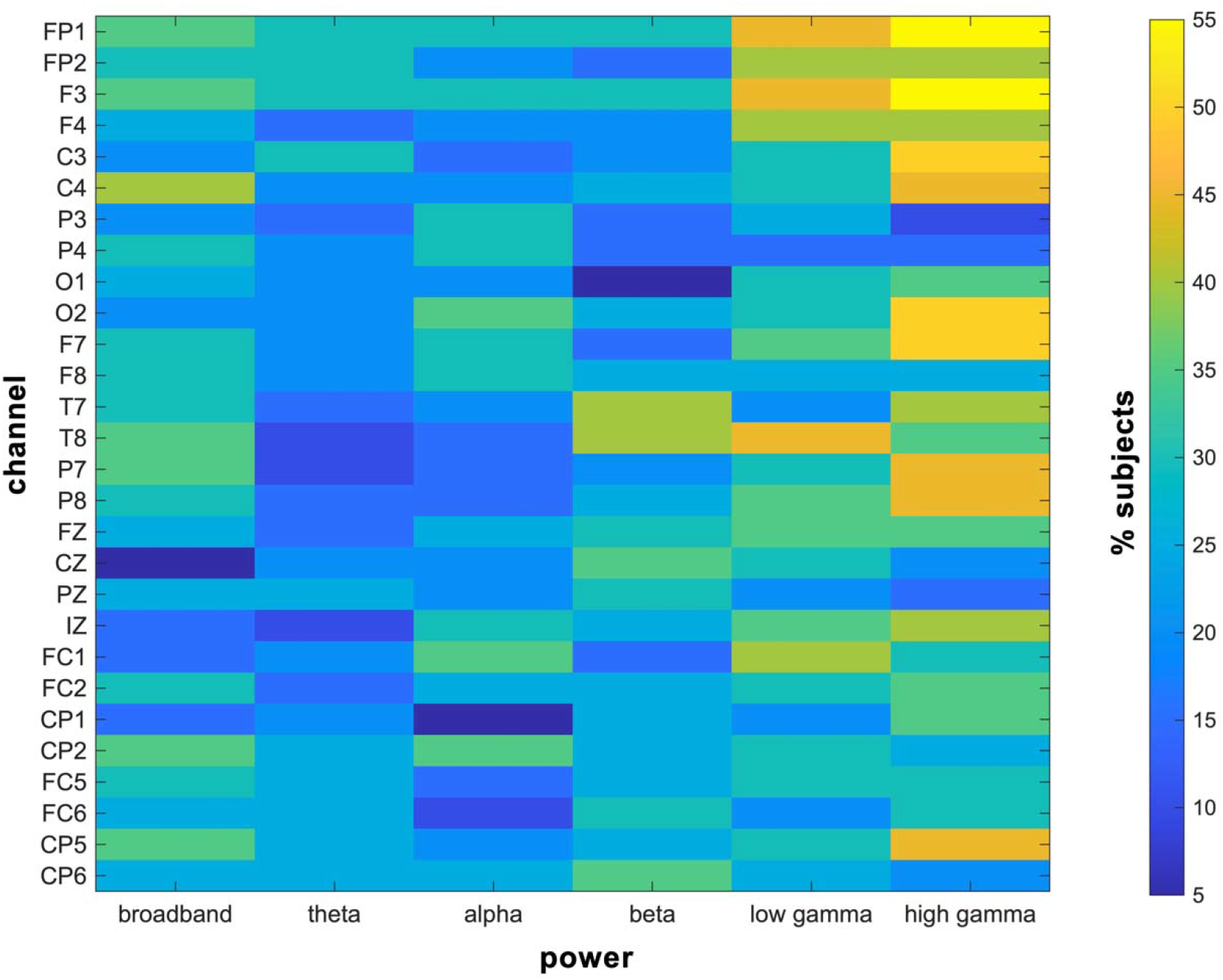
Power spectral features commonly included in each participant’s optimized classifier. Gamma power features across the scalp were included in approximately half of all participants’ final classifiers.

Given that the features used to successfully discriminate between high and low M1 excitability states differed across participants, we also examined cross-participant generalization of each participant’s optimized classifier. Specifically, we tested whether personalized classifiers trained using power spectral features calculated from EEG signals immediately preceding TMS (0.505 to 0.005 s before each TMS pulse) could discriminate between high and low M1 excitability states in all other participants. Personalized classifiers trained using all trials from 5 of the 20 individual participants performed above theoretical chance when tested on all other participants (p < 0.05, Figure 4a) while this was not the case for the remaining 15 (p > 0.07). This corresponded to group-level generalization performance AUC values of 0.51 ± 0.05 (range = 0.33 – 0.66, Figure 4b), which did not exceed theoretical chance (p = 0.07). Generalization performance was also significantly worse than personalized participant-specific classification performance (p = 4.78 × 10^-5^, see also Figure 2a).

**Figure 4.**
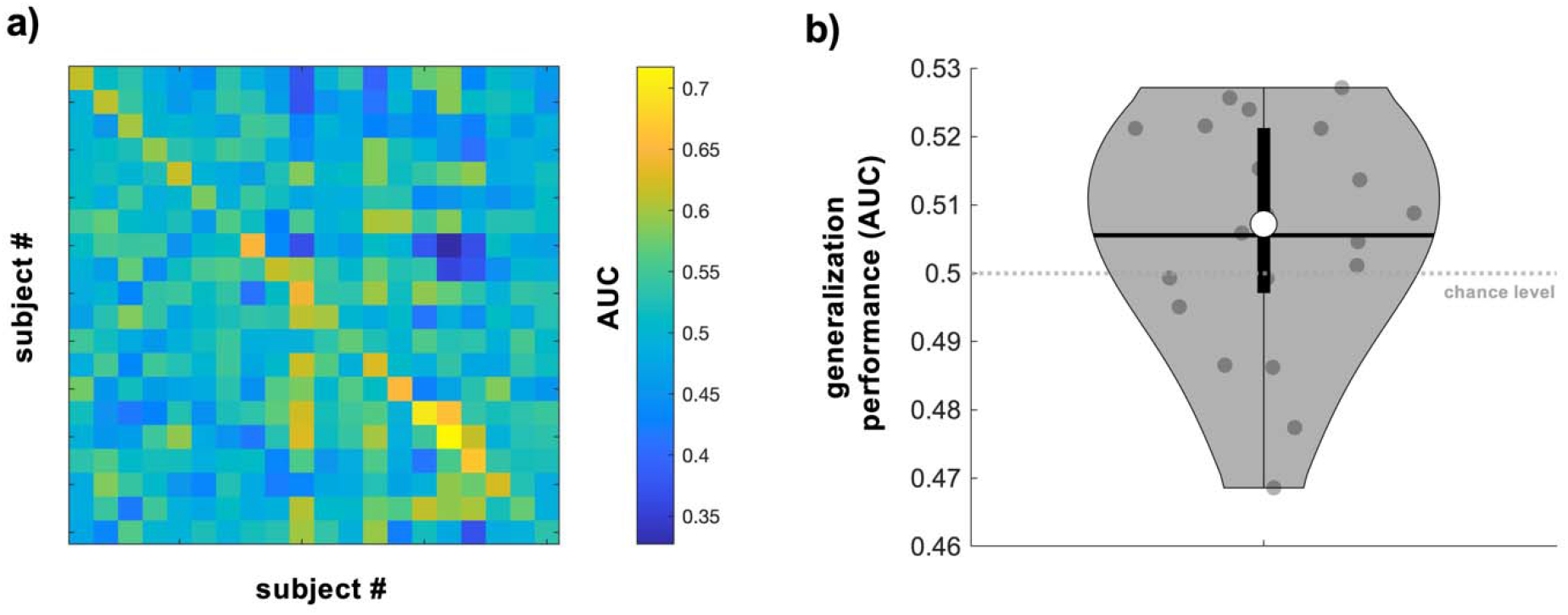
Cross-participant generalization of personalized classifiers. a) Matrix of AUC values reflecting cross-participant generalization performance of classifiers trained using power spectral features obtained from EEG signals immediately preceding TMS (0.505 – 0.005 s before TMS, classifiers were trained using all trials per participant). Warmer and cooler colors indicate higher and lower AUC values, respectively. AUC values along the diagonal indicate performance of personalized classifiers and thus are identical to those shown in Figure 2a. b) AUC values indicating generalization performance averaged across all testing participants per training participant. AUC values exceeded theoretical chance in 5 of the 20 participants but not at the group level. For b, grey dots = individual participant AUC values, thick whisker = interquartile range, thin whisker = full range, white dot = group median, black horizontal line = group mean, grey dashed line = theoretical chance level.

We next evaluated the temporal evolution of classification performance over three seconds leading up to TMS delivery. Personalized classifiers trained and tested on power spectral features extracted from EEG data within each pre-TMS window (3.005 –2.505 s, 2.505 – 2.005 s, 2.005 – 1.505 s, 1.505 – 1.005 s, 1.005 – 0.505 s before TMS) showed above-chance performance for all time windows (empirical chance = 0.55, Figure 5a, p < 1.36 × 10^-4^ for all). Classifier performance immediately before TMS delivery did not differ from performance for any other time windows preceding TMS delivery (p > 0.08 for all). Further, AUC values of personalized classifiers trained using features obtained immediately before TMS delivery (0.505 – 0.005 s before TMS) and tested on each preceding time window also exceeded empirical chance for all time windows (p < 1.18 × 10^-4^ for all; Figure 5b). Classification AUC values obtained using features calculated from EEG data immediately preceding TMS delivery (0.505 – 0.005 s before TMS) did not differ from generalization AUC values obtained from any other time windows preceding TMS (p > 0.06 for all).

**Figure 5.**
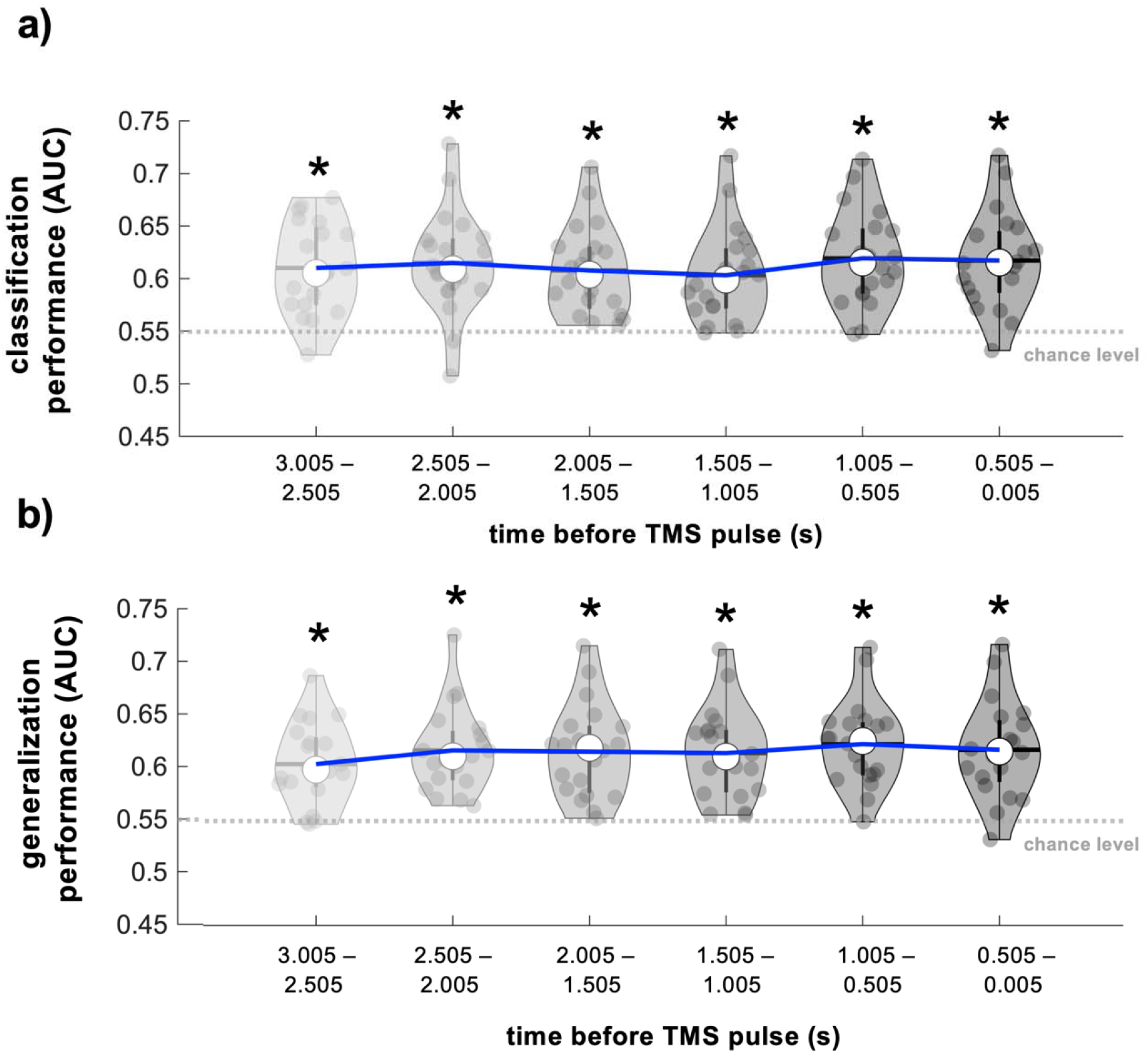
Evaluation of personalized classification performance over time. a) AUC values reflecting performance of classifiers trained and tested using power spectral features obtained from EEG data in different time windows preceding TMS delivery. Classification AUC values for the time window closest to TMS delivery (0.500 – 0.005 s before TMS) did not differ from classification AUC values obtained for all other time windows. b) AUC values reflecting the extent to which personalized classifiers trained using power spectral features obtained from EEG data immediately before TMS delivery (0.500 – 0.005 s, far right) generalized to earlier time windows. Classification AUC values for the time window closest to TMS delivery (0.500 – 0.005 s before TMS) did not differ from generalization AUC values for all other time windows. Note that for both panels, the data depicted on the right-most violin plot reflects the same classification AUC values depicted in Figure 2a. Time windows used to calculate power spectral features are depicted on the x axes. Blue lines indicate group-level AUC values approaching TMS delivery. Grey dots represent individual participant AUC values, thick whisker represents interquartile range, thin whisker represents full range, white dot represents the group median and grey/black horizontal lines represent the group mean. The grey dashed lines indicate the empirical chance level. * Indicates statistical significance of AUC values when compared to empirical chance.

We also evaluated the number of trials needed to achieve successful classification by training and testing personalized classifiers using varying dataset sizes (50 to 450 trials in 50 trial increments). Classifier performance exceeded empirical chance for all dataset sizes (chance = 0.55, p < 1.02 × 10^-4^) but was highest when using datasets containing 50 trials (AUC = 0.72 ± 0.01, range = 0.62 – 0.87, Figure 6a). Consistent with this, AUC values obtained from classifiers built using 50 trials were significantly higher than AUC values obtained using classifiers built using 150, 200, 250, 300, 350, 400, or 450 trials (p < 0.03). AUC values obtained from classifiers built using 50 trials did not differ from AUC values obtained from classifiers built using 100 trials (p = 0.09). Further, classifier-predicted MEP amplitude modulation significantly exceeded 1 for all dataset sizes (p < 1.36 × 10^-4^), indicating that classifier-predicted high MEP amplitudes were larger than classifier-predicted low MEP amplitudes for each dataset size (Figure 6b). However, classifier-predicted MEP amplitude modulation obtained from classifiers built using 50 trials did not differ from classifier-predicted MEP amplitude modulation obtained from classifiers built using any other dataset size (p = 0.37).

**Figure 6.**
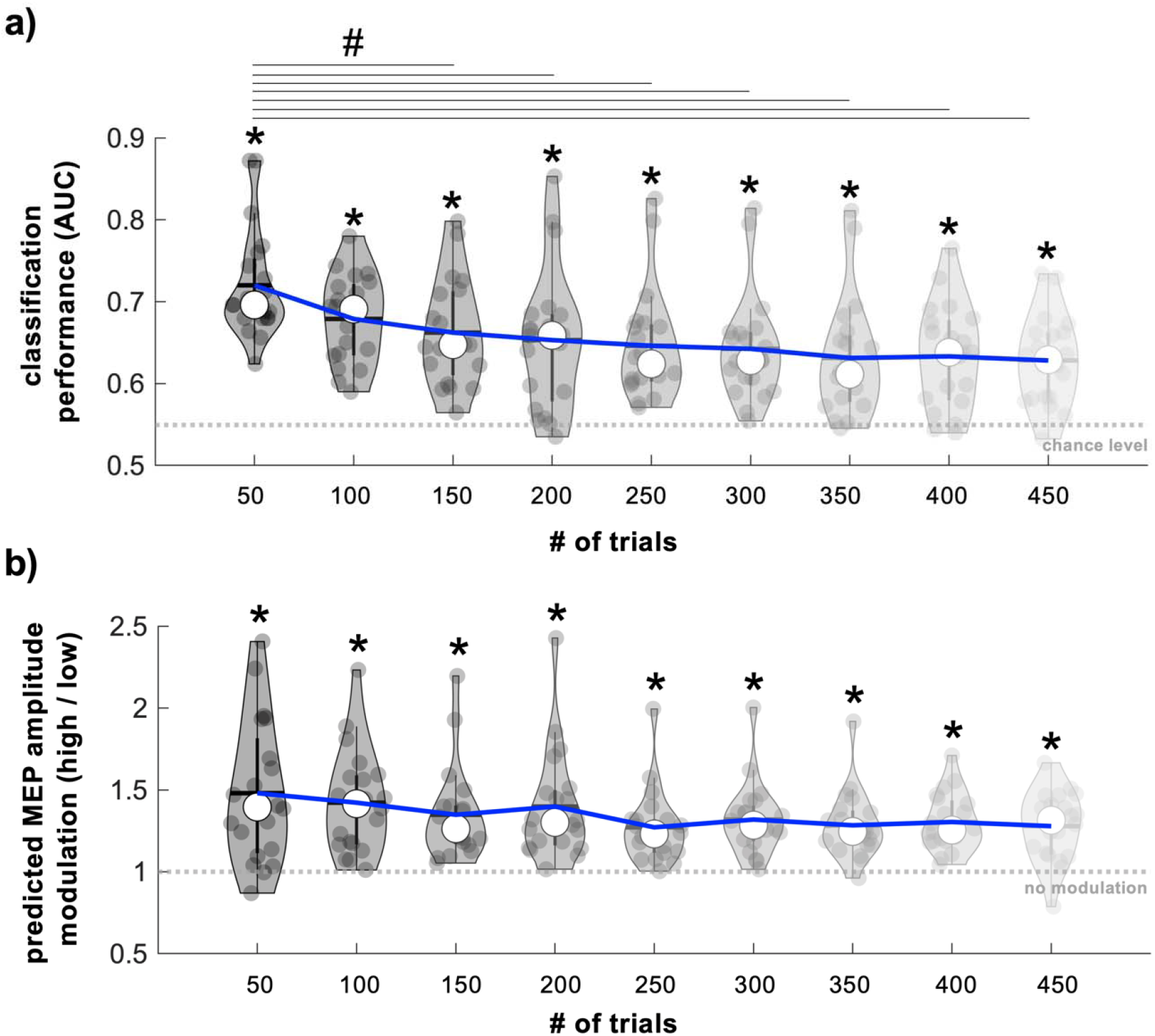
Influence of trial number on classifier performance and classifier-predicted MEP amplitude modulation. a) AUC values reflecting performance of classifiers trained and tested using varying dataset sizes. AUC values exceeded empirical chance for all dataset sizes. However, AUC values obtained from classifiers built using 50 trials were significantly larger than AUC values obtained from classifiers built using 150, 200, 250, 300, 350, 400, or 450 trials. The grey dashed line indicates empirical chance level. * Indicates statistical significance of AUC values when compared to empirical chance. # Indicates significant pairwise comparisons. b) Classifier-predicted MEP amplitude modulation obtained from classifiers built using varying dataset sizes. Predicted MEP amplitude modulation (classifier-predicted high MEP amplitudes / classifier-predicted low MEP amplitudes) was larger than 1 for all dataset sizes and did not differ across dataset sizes. The grey dashed line indicates no MEP amplitude modulation. * Indicates statistical significance of predicted MEP amplitude values when compared to 1. For both panels, blue lines indicate group means for each dataset size. Grey dots represent individual participant values, thick whisker represent interquartile range, thin whisker represent full range, white dots represent the group median and grey/black horizontal lines represent group mean.

To assess our approach’s potential usefulness in a real-time setting, we also calculated the time needed to compute power spectral features and predict the M1 excitability state. At the group level, this took 9.6 ± 3.98 × 10^-5^ ms (range = 9.5 – 10.4 ms). The number of features used for each participant’s final classifier and that participant’s mean computational time were not significantly correlated (rho = 0.15, p = 0.26), suggesting that the computational time needed for our approach does not depend on the number of features incorporated into each participant’s personalized classifier.

## 4. Discussion

In this study, we used multivariate pattern classification to identify personalized M1 excitability states from human EEG recordings. We found that personalized classifiers successfully discriminated between low and high M1 excitability states in 80% of tested participants. MEPs evoked during classifier-predicted high M1 excitability states were ~24% larger than those evoked during classifier-predicted low M1 excitability states, and personalized classifiers did not generalize across participants. Further, classifiers trained using features extracted from EEG data up to 3 seconds before TMS delivery performed above chance. Overall, our findings demonstrate that individual participants exhibit unique brain activity patterns that successfully predict M1 excitability states and that these patterns can be efficiently captured using simple power spectral features obtained from low-resolution EEG recordings.

Historically, TMS has been delivered to the brain irrespective of ongoing brain activity. Early studies showed that TMS protocols using different stimulation rates and burst patterns induce either LTP- or LTD-like plasticity within human M1.^11–14^ However, more recent reports suggest that the plastic effects of TMS interventions may depend more strongly on the excitability state at the time of stimulation than the specific pulse rate or pattern,^20,21^ consistent with findings in animal models.^39–42^ For example, whether hippocampal stimulation induces LTP or LTD critically depends on the theta phase (which is closely tied to hippocampal excitability^43^) at the exact moment of stimulation.^39–41^ Accumulating evidence thus suggests that TMS protocols are most likely to induce LTP- and LTD-like plasticity when delivered during high and low excitability states, respectively, motivating development of effective and generalizable methods capable of predicting the presence of such states.^18–20,44^ Along these lines, current state-of-the-art methods use oscillatory properties that correlate with excitability at the group level.^20–22,26,29^ However, adapting these methods to individual participants can be challenging,^28^ and properly applying them in patients with brain lesions or other forms of structural brain damage raises additional issues.^29^ Here, we introduce a straightforward, personalized decoding-based analytical framework that successfully predicts the occurrence of high and low M1 excitability states in individual participants.

The decoding-based approach described here holds several advantages over other recent brain state detection algorithms.^20–22,26,45^ First, our approach is entirely personalized and thus does not require *a priori* knowledge of the relationship between cortical excitability and EEG activity. Instead, it discovers patterns of brain activity that correspond to high and low M1 excitability states using supervised machine learning at the individual participant level. Compared to “top down” hypothesis-driven studies that investigate the relationship between pre-defined brain states and M1 excitability, our data-driven “bottom up” approach increases the likelihood of successfully detecting high and low excitability states. Second, because it is fully personalized, our decoding-based approach in principle does not require any specific cortical structure. This characteristic makes it well-suited for brain state-dependent TMS in patients with brain damage that can alter the relationship between neuronal activity and scalp-level EEG signals (i.e., cortical stroke; see^29,46^). Third, our classification approach requires little EEG preprocessing. Personalized classifiers built using highly preprocessed EEG signals^45^ are not likely to generalize well to noisy real-world EEG signals regardless of their performance during cross-validation. For this reason, we intentionally did not include time-intensive analytical steps like independent components analysis in our preprocessing approach. We therefore expect our method to perform well in a real-time setting. A fourth advantage is that we used all available trials to build personalized classifiers. In contrast, a recent study built personalized classifiers using only those trials resulting in the most extreme MEP amplitudes (i.e., the largest and smallest^45^), reporting classification accuracy of ~67%. Although retaining only those trials associated with extreme MEP amplitudes improves performance during cross-validation, classifiers trained in this way may not generalize well to a real-time setting where brain states outside of these extremes are common. Consistent with this, we found that retaining only those trials associated with extreme MEP amplitudes in our dataset artificially boosted classifier performance (AUC = 0.67 ± 0.01, range = 0.57 – 0.82).

In addition to successfully discriminating between low and high M1 excitability states, the brain states identified here lasted for seconds at a time. Specifically, our results showed that EEG data obtained up to 3 seconds before TMS delivery successfully discriminated between low and high M1 excitability states, yet our computational approach only requires ~10 ms to predict the upcoming M1 excitability state. These results have clear practical implications for brain state-dependent TMS: because the time needed to predict the upcoming M1 excitability state is substantially lower than the duration of M1 excitability states themselves, TMS could in the future simply be triggered upon the detection of the desired excitability state. This triggering-based approach contrasts starkly with computationally-intensive phase-dependent TMS algorithms that use autoregressive forward prediction to estimate rapidly changing instantaneous oscillatory phases at some future time point^20–22,26,29,44^ and are much easier to implement (see^47^). Further, the MEP amplitude modulation between classifier-predicted high and low M1 excitability states seen here (~24%) closely matches that observed when using phase-dependent M1 TMS,^20,21,48,49^ suggesting similar efficacy despite a reduced computational load.

Personalized classifiers did not generalize across participants and classification at the individual participant level significantly outperformed cross-participant generalization, indicating the importance of personalized EEG features when predicting M1 excitability levels. This notion is further supported by the fact that the features included in each participant’s final classifier varied across participants. Previous work showed that personalized features can explain up to half of the variance in time-varying brain activity^50^ and can reliably identify individual participants.^51^ Given these prior reports, it is perhaps not surprising that personalized brain activity patterns also predict M1 excitability states. Although no single feature was consistently included in all participant’s final classifiers, gamma-range features across the scalp were included in approximately half of participants’ final classifiers (see Figure 3, yellow and orange cells). Given that gamma rhythms are closely tied to neuronal activation^52^ and attention,^53^ the gamma-range features identified here may capture spontaneous fluctuations in attention or arousal. Consistent with this possibility, recent work has shown that M1 excitability is dynamically shaped by attention and alertness^54,55^. Still, whether the brain states identified here are stable across multiple recording sessions has yet to be established. Determining whether personalized classifiers successfully generalize across multiple sessions will be important for multi-session decoding-based brain state-dependent neuromodulation interventions.

The current study used several hundred trials to build personalized classifiers. However, collecting a large TMS-EEG-EMG dataset and using it to build a personalized classifier before delivering a brain state-dependent TMS intervention may not always be feasible. We therefore evaluated whether our approach could discriminate between low and high M1 excitability states using datasets of varying sizes. This analysis revealed that personalized classifiers performed best when using only 50 trials, suggesting that 50 trials may be adequate to successfully discriminate between low and high M1 excitability states. One possible explanation for the observed improvement in performance when using datasets with fewer trials is that participants became fatigued during the long experimental session. Changes in alertness may have modified the pattern of brain activity that best predicted low and high M1 excitability states leading to poorer overall performance as more trials were included. Conventional braincomputer interfaces often encounter a similar problem when moving from an offline training session to an online control session.^56^ Adaptive methods that iteratively update each participant’s personalized classifier could feasibly compensate for any fatigue-related reductions in classifier performance during brain state-dependent TMS interventions^56,57^.

Although the personalized classifiers used here performed above chance in the majority of tested participants, their overall performance is lower than classifiers used for other real-time EEG analysis applications like brain-computer interfaces.^58^ However, despite the modest performance levels achieved here, classifier-predicted high and low M1 excitability states were associated with an ~24% difference in MEP amplitudes. Previous brain state-dependent TMS studies showed that applying TMS during high excitability brain states during which TMS evokes MEPs that are ~14-36% larger than those evoked during low excitability states substantially enhances the neuroplastic effects of TMS interventions.^20–22^ Based on these findings, we expect that translating our approach to a real-time setting will be sufficient to strengthen the effects of M1 TMS interventions. In addition, the fact that MEPs elicited during true high M1 excitability states were ~260% larger than those elicited during true low M1 excitability states indicates that further improving classifier performance should produce even larger MEP amplitude differences between classifier-predicted low and high M1 excitability states. Future work should focus on optimizing classifier performance to maximize the beneficial effects of personalized brain state-dependent TMS.

In conclusion, we report that individual participants exhibit unique brain activity patterns that predict M1 excitability levels. These patterns were successfully decoded in 80% of tested participants using only simple power spectral features obtained from low-density scalp EEG signals. MEP amplitudes elicited during classifier-predicted high M1 excitability states were also ~24% larger than those elicited during classifier-predicted low M1 excitability states. Finally, these brain states lasted for seconds at a time, making them ideal targets for efficient brain state-dependent TMS. Overall, the approach presented here constitutes a key step towards efficient, flexible, and fully personalized decoding-based brain state-dependent TMS in the future.

## Acknowledgements

Data included in this publication were acquired at the Human Cortical Physiology and Neurorehabilitation Section (directed by Leonardo Cohen) at NINDS and are publicly available. This research did not receive any specific grant from funding agencies in the public, commercial, or not-for-profit sectors.

## Author contributions

SJH: conceptualization, methodology, software, validation, formal analysis, investigation, data curation, writing, visualization.

RQ: conceptualization, methodology, validation, formal analysis, writing.

## Data availability

All data are publicly available at https://openneuro.org/datasets/ds002094/versions/1.0.0.

## References

1. George, M. S. et al. Daily repetitive transcranial magnetic stimulation (rTMS) improves mood in depression. NeuroReport 6, 1853–1856 (1995).

2. O’Reardon, J. P. et al. Efficacy and Safety of Transcranial Magnetic Stimulation in the Acute Treatment of Major Depression: A Multisite Randomized Controlled Trial. Biological Psychiatry 62, 1208–1216 (2007).

3. Gorelick, D. A., Zangen, A. & George, M. S. Transcranial magnetic stimulation (TMS) in the treatment of substance addiction. 1327, 79–93 (2014).

4. Hanlon, C. A., Dowdle, L. T. & Scott Henderson, J. Modulating Neural Circuits with Transcranial Magnetic Stimulation: Implications for Addiction Treatment Development. Pharmacological Reviews 70, 661–683 (2018).

5. Ekhtiari, H. et al. Transcranial electrical and magnetic stimulation (tES and TMS) for addiction medicine: A consensus paper on the present state of the science and the road ahead. Neuroscience & Biobehavioral Reviews 104, 118–140 (2019).

6. Wang, J. X. et al. Memory Enhancement: Targeted enhancement of cortical-hippocampal brain networks and associative memory. Science 345, 1054–1057 (2014).

7. Nilakantan, A. S. et al. Network-targeted stimulation engages neurobehavioral hallmarks of age-related memory decline. Neurology 92, E2349–E2354 (2019).

8. Tazoe, T. & Perez, M. A. Effects of Repetitive Transcranial Magnetic Stimulation on Recovery of Function After Spinal Cord Injury. Archives of Physical Medicine and Rehabilitation 96, S145–S155 (2015).

9. Hummel, F. C. & Cohen, L. G. Non-invasive brain stimulation: a new strategy to improve neurorehabilitation after stroke? The Lancet Neurology 5, 708–712 (2006).

10. Webster, B. R., Celnik, P. A. & Cohen, L. G. Noninvasive Brain Stimulation in Stroke Rehabilitation. NeuroRX 3, 474–481 (2006).

11. Pascual-leone, A., Valls-solé, J., Wassermann, E. M. & Hallett, M. Responses to rapid-rate transcranial magnetic stimulation of the human motor cortex. Brain 117, 847–858 (1994).

12. Chen, R. et al. Depression of motor cortex excitability by low-frequency transcranial magnetic stimulation. Neurology 48, 1398–1403 (1997).

13. Stefan, K., Kunesch, E., Cohen, L. G., Benecke, R. & Classen, J. Induction of plasticity in the human motor cortex by paired associative stimulation. Brain 123, 572–584 (2000).

14. Huang, Y. Z., Edwards, M. J., Rounis, E., Bhatia, K. P. & Rothwell, J. C. Theta burst stimulation of the human motor cortex. Neuron 45, 201–206 (2005).

15. Ziemann, U. & Siebner, H. R. Inter-subject and Inter-session Variability of Plasticity Induction by Non-invasive Brain Stimulation: Boon or Bane? Brain stimulation 8, 662–663 (2015).

16. Goldsworthy, M. R., Hordacre, B., Rothwell, J. C. & Ridding, M. C. Effects of rTMS on the brain: is there value in variability? Cortex 139, 43–59 (2021).

17. López-Alonso, V., Cheeran, B., Río-Rodríguez, D. & Fernández-Del-Olmo, M. Interindividual Variability in Response to Non-invasive Brain Stimulation Paradigms. Brain Stimulation 7, 372–380 (2014).

18. Kraus, D. et al. Recruitment of additional corticospinal pathways in the human brain with state-dependent paired associative stimulation. Journal of Neuroscience 38, 1396–1407 (2018).

19. Kraus, D. et al. Brain–robot interface driven plasticity: Distributed modulation of corticospinal excitability. NeuroImage 125, 522–532 (2016).

20. Zrenner, C., Desideri, D., Belardinelli, P. & Ziemann, U. Real-time EEG-defined excitability states determine efficacy of TMS-induced plasticity in human motor cortex. Brain Stimulation 11, 374–389 (2018).

21. Baur, D. et al. Induction of LTD-like corticospinal plasticity by low-frequency rTMS depends on pre-stimulus phase of sensorimotor μ-rhythm. Brain Stimulation 13, 1580–1587 (2020).

22. Hussain, S. J. et al. Phase-dependent offline enhancement of human motor memory. Brain Stimulation 14, 873–883 (2021).

23. Haegens, S., Nácher, V., Luna, R., Romo, R. & Jensen, O. α-Oscillations in the monkey sensorimotor network influence discrimination performance by rhythmical inhibition of neuronal spiking. Proceedings of the National Academy of Sciences of the United States of America 108, 19377–19382 (2011).

24. Berger, B., Minarik, T., Liuzzi, G., Hummel, F. C. & Sauseng, P. EEG oscillatory phase-dependent markers of corticospinal excitability in the resting brain. BioMed Research International 2014, (2014).

25. Hussain, S. J. et al. Sensorimotor oscillatory phase-power interaction gates resting human corticospinal output. Cerebral Cortex 29, 3766–3777 (2019).

26. Madsen, K. H. et al. No trace of phase: Corticomotor excitability is not tuned by phase of pericentral mu-rhythm. Brain Stimulation 12, 1261–1270 (2019).

27. Bergmann, T. O., Lieb, A., Zrenner, C. & Ziemann, U. Pulsed facilitation of corticospinal excitability by the sensorimotor μ-alpha rhythm. Journal of Neuroscience 39, 10034–10043 (2019).

28. Zrenner, C. et al. The shaky ground truth of real-time phase estimation. NeuroImage 214, 116761 (2020).

29. Hussain, S. J. et al. Phase-dependent transcranial magnetic stimulation of the lesioned hemisphere is accurate after stroke. Brain Stimulation 13, 1354–1357 (2020).

30. Hussain, S. J., Cohen, L. G. & Bönstrup, M. Beta rhythm events predict corticospinal motor output. Scientific Reports 9, 1–10 (2019).

31. Awiszus F, B. J. TMS motor threshold assessment tool (MTAT 2.0). Brain Stimulation Laboratory, Medical University of South Carolina, USA. (2011).

32. Oostenveld, R., Fries, P., Maris, E. & Schoffelen, J. M. FieldTrip: Open source software for advanced analysis of MEG, EEG, and invasive electrophysiological data. Computational Intelligence and Neuroscience 2011, (2011).

33. Farzan, F. et al. Reliability of long-interval cortical inhibition in healthy human subjects: A TMS-EEG study. Journal of Neurophysiology 104, 1339–1346 (2010).

34. Jensen, O. et al. On the human sensorimotor-cortex beta rhythm: Sources and modeling. NeuroImage 26, 347–355 (2005).

35. Nowak, M., Zich, C. & Stagg, C. J. Motor Cortical Gamma Oscillations: What Have We Learnt and Where Are We Headed? Current Behavioral Neuroscience Reports 2018 5:2 5, 136–142 (2018).

36. Treder, M. S. MVPA-Light: A Classification and Regression Toolbox for Multi-Dimensional Data. Frontiers in Neuroscience 14, 289 (2020).

37. Benjamini, Y. & Hochberg, Y. Controlling the False Discovery Rate: A Practical and Powerful Approach to Multiple Testing. Journal of the Royal Statistical Society: Series B (Methodological) 57, 289–300 (1995).

38. King, J. R. & Dehaene, S. Characterizing the dynamics of mental representations: the temporal generalization method. Trends in Cognitive Sciences 18, 203–210 (2014).

39. Pavlides, C., Greenstein, Y. J., Grudman, M. & Winson, J. Long-term potentiation in the dentate gyrus is induced preferentially on the positive phase of θ-rhythm. Brain Research 439, 383–387 (1988).

40. Huerta, P. T. & Lisman, J. E. Heightened synaptic plasticity of hippocampal CA1 neurons during a Cholinergically induced rhythmic state. Nature 1993 364:6439 364, 723–725 (1993).

41. Huerta, P. T. & Lisman, J. E. Bidirectional synaptic plasticity induced by a single burst during cholinergic theta oscillation in CA1 in vitro. Neuron 15, 1053–1063 (1995).

42. Zanos, S., Rembado, I., Chen, D. & Fetz, E. E. Phase-Locked Stimulation during Cortical Beta Oscillations Produces Bidirectional Synaptic Plasticity in Awake Monkeys. Current Biology 28, 2515–2526.e4 (2018).

43. Buzsáki, G. Theta Oscillations in the Hippocampus. Neuron 33, 325–340 (2002).

44. Shirinpour, S., Alekseichuk, I., Mantell, K. & Opitz, A. Experimental evaluation of methods for real-time EEG phase-specific transcranial magnetic stimulation. Journal of Neural Engineering 17, (2020).

45. Metsomaa, J., Belardinelli, P., Ermolova, M., Ziemann, U. & Zrenner, C. Causal decoding of individual cortical excitability states. NeuroImage 245, 118652 (2021).

46. Baumann, S. B., Wozny, D. R., Kelly, S. K. & Meno, F. M. The electrical conductivity of human cerebrospinal fluid at body temperature. IEEE Transactions on Biomedical Engineering 44, 220–223 (1997).

47. Bergmann, T. O. et al. EEG-Guided Transcranial Magnetic Stimulation Reveals Rapid Shifts in Motor Cortical Excitability during the Human Sleep Slow Oscillation. Journal of Neuroscience 32, 243–253 (2012).

48. Hussain, S. J. et al. Sensorimotor Oscillatory Phase-Power Interaction Gates Resting Human Corticospinal Output. doi:10.1093/cercor/bhy255.

49. Bergmann, T. O., Lieb, A., Zrenner, C. & Ziemann, U. Pulsed Facilitation of Corticospinal Excitability by the Sensorimotor μ-Alpha Rhythm. Journal of Neuroscience 39, 10034–10043 (2019).

50. Gratton, C. et al. Functional Brain Networks Are Dominated by Stable Group and Individual Factors, Not Cognitive or Daily Variation. Neuron 98, 439–452.e5 (2018).

51. Finn, E. S. et al. Functional connectome fingerprinting: identifying individuals using patterns of brain connectivity. Nature Neuroscience 2015 18:11 18, 1664–1671 (2015).

52. Niessing, J. et al. Neuroscience: Hemodynamic signals correlate tightly with synchronized gamma oscillations. Science 309, 948–951 (2005).

53. Fries, P., Reynolds, J. H., Rorie, A. E. & Desimone, R. Modulation of oscillatory neuronal synchronization by selective visual attention. Science 291, 1560–1563 (2001).

54. Rösler KM, Etter C, Truffert A, Hess CW & Magistris. Rapid cortical motor output map changes assessed by the triple stimulation technique. NeuroReport 10, 579–583 (1999).

55. Noreika, V. et al. Alertness fluctuations when performing a task modulate cortical evoked responses to transcranial magnetic stimulation. NeuroImage 223, 117305 (2020).

56. Shenoy, P., Krauledat, M., Blankertz, B., Rao, R. P. N. & Müller, K. R. Towards adaptive classification for BCI*. Journal of Neural Engineering 3, R13 (2006).

57. Vidaurre, C., Schlöogl, A., Cabeza, R., Scherer, R. & Pfurtscheller, G. A fully on-line adaptive BCI. IEEE Transactions on Biomedical Engineering 53, 1214–1219 (2006).

58. Blankertz, B., Curio, G. & Müller, K.-R. Classifying Single Trial EEG: Towards Brain Computer Interfacing.

